# Mendelian randomization: a novel approach for the prediction of adverse drug events and drug repurposing opportunities

**DOI:** 10.1101/105338

**Authors:** Venexia M Walker, George Davey Smith, Neil M Davies, Richard M Martin

**Affiliations:** School of Social and Community Medicine, University of Bristol, United Kingdom; MRC University of Bristol Integrative Epidemiology Unit, United Kingdom

**Keywords:** Mendelian randomization, pharmacovigilance, drug repurposing, adverse drug events

## Abstract

Identification of unintended drug effects, specifically drug repurposing opportunities and adverse drug events, maximizes the benefit of a drug and protects the health of patients. However, current observational research methods are subject to several biases. These include confounding by indication, reverse causality, and missing data. We propose that Mendelian randomization (MR) offers a novel approach for the prediction of unintended drug effects. In particular, we advocate the synthesis of evidence from this method and other approaches, in the spirit of triangulation, to improve causal inferences concerning drug effects. MR overcomes some of the limitations associated with the existing methods in this field. Furthermore, it can be applied either pre- or post-approval of the drug and could therefore prevent the potentially harmful exposure of patients in clinical trials and beyond. The potential of MR as a pharmacovigilance and drug repurposing tool is yet to be realized and could both help prevent adverse drug events and identify novel indications for existing drugs in the future.

## KEY MESSAGES

- We propose that the prediction of unintended drug effects using Mendelian randomization can overcome some of the limitations associated with existing methods; including confounding by indication, reverse causality, and missing data.
- We demonstrate the potential of Mendelian randomization for predicting unintended drug effects using examples from the literature of studies that have assessed recognized unintended drug effects.
- We advocate the synthesis of evidence from Mendelian randomization and other approaches, in the spirit of triangulation, to improve causal inferences concerning drug effects.

Adverse drug events and drug repurposing opportunities are both unintended drug effects. Drug repurposing, defined as the application of known drugs to new indications (1), offers a time and cost effective alternative to traditional drug development. Adverse drug events, defined as any unwanted reaction to a drug (2), risk patient safety and increase the burden on healthcare systems. The opportunities offered by drug repurposing and the potential harm caused by adverse drug events means the identification of unintended drug effects is necessary to maximize the benefit of a drug and protect health.

Unintended effects of drugs can be discovered throughout the drug development process. However, prior to the approval of a novel drug, its risk-benefit profile cannot be fully known. This is because pre-approval clinical trials are principally for demonstrating the drug’s efficacy for its intended indication. This limits the trial’s ability to assess safety and identify novel indications in a number of ways. (3) Firstly, the comparatively small number of patients exposed to a drug during a pre-approval clinical trial means that only very common or very large drug effects can be detected. Secondly, the length of time that patients are exposed to the drug in this setting is relatively short. Thirdly, the recorded data may not include the necessary information to identify previously unknown drug effects or those that are unrelated to the drug’s indication. Finally, the participants of a study may not represent the broad range of patients seen in clinical practice. As a result of this, continued assessment of drugs post-approval is necessary in order to fully develop their profile.

Post-approval of a drug, unintended drug effects can be identified in a number of ways. Adverse drug events are primarily identified through the use of spontaneous reporting systems (4–6), which rely on healthcare professionals and members of the public to report suspected drug effects. Drug repurposing opportunities are often sought directly by pharmaceutical companies using purpose built drug repurposing technology platforms, due to their desirable risk-versus-reward trade off. (7) Strong signals from these databases and technology platforms are then investigated using data from a range of sources, including: randomized clinical trials either pre- or post-approval of the drug, meta-analyses of such trials, observational studies and information from basic science. (3) However these methods, particularly spontaneous reporting systems, suffer from several biases including their inability to determine causality, over reporting from media coverage, confounding by indication and other usually unobserved confounders. Minimizing these biases is therefore key to determining which of these signals indicate a true unintended drug effect.

We propose that Mendelian randomization (MR) (8–10) offers a novel approach for the prediction of unintended drug effects, which overcomes some of the limitations associated with existing methods. MR assesses the causal effect of an exposure on an outcome by using a genetic variant as a proxy for exposure. The principle behind this is that randomization occurs naturally at conception when genetic variants are allocated at random to individuals from their parents. The genetic variants allocated at this time are part of the germline genome – this is represented in studies by the data collected from genotyping. The alternative to the germline genome is the somatic genome – this describes the genetic variants that can alter post-conception. Using germline genetic variants for the prediction of unintended drug effects limits bias due to non-genetic confounding, particularly confounding by indication, and reverse causation because these variants cannot be influenced by environmental, lifestyle or disease-related factors operating later in life. For example, consider statins prescribed for the prevention of coronary heart disease (CHD). Statins inhibit the enzyme 3-hydroxy-3-methylglutaryl-CoA reductase (HMGCR) to lower low density lipoprotein (LDL) cholesterol and consequently reduce the risk of CHD. To investigate unintended drug effects associated with statins, a MR study would use a single-nucleotide polymorphism (SNP) located on the HMGCR gene as a proxy for exposure to statins. The application of MR, as described here, is presented in Figure 1 and later discussed in the context of a study. (11) It is worth noting at this point that MR can either be applied before other studies are launched to generate information about potential unintended drug effects, or after other studies are completed to strengthen the evidence provided by them. In particular, we advocate the synthesis of evidence from MR and other approaches, in the spirit of triangulation, to improve causal inferences concerning drug effects. (12) We also encourage the use of MR to enhance existing techniques, such as some in silico approaches, which can use predicted genes in their analysis. (13)

**Figure 1:**
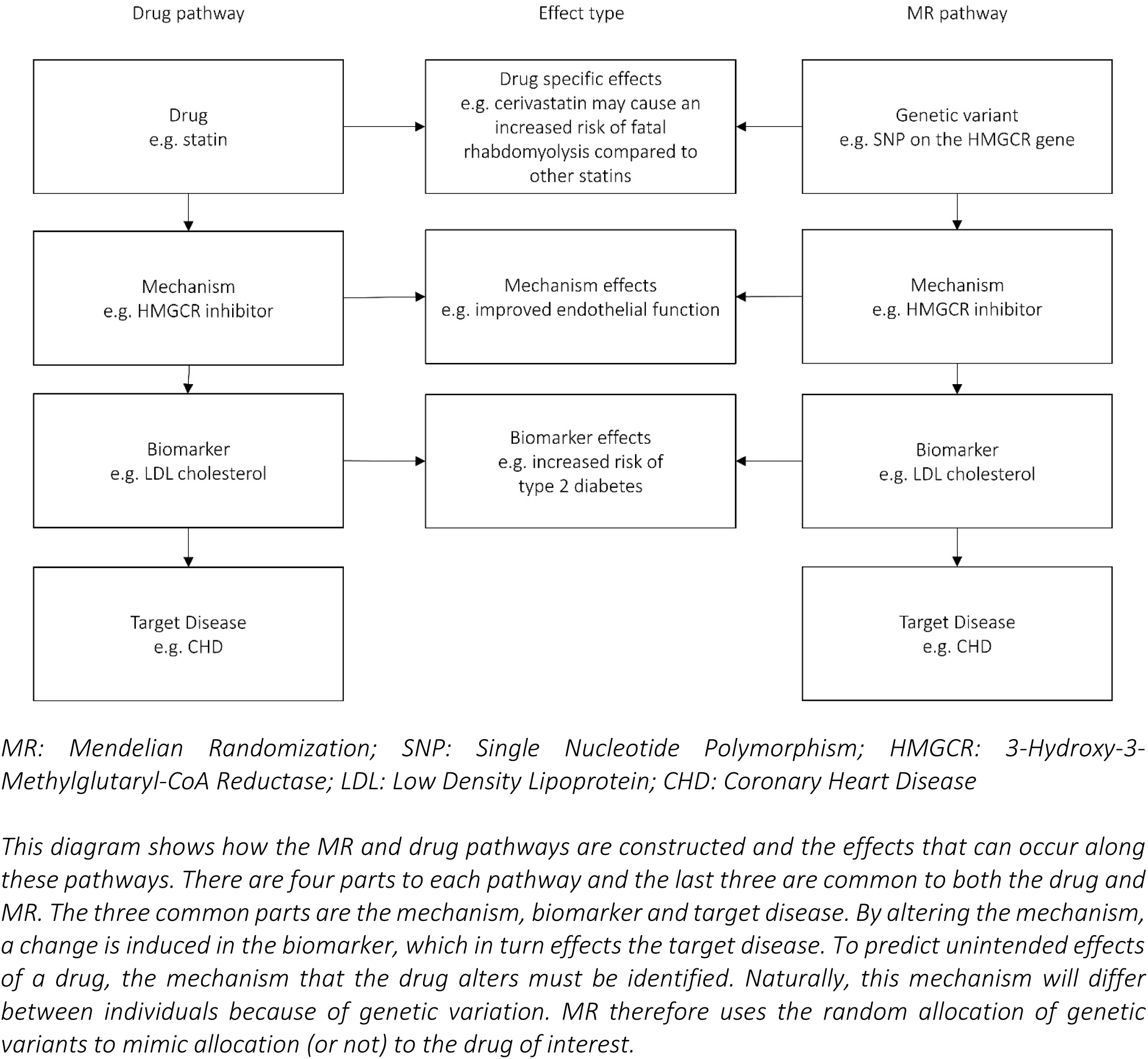
The process by which MR mimics the action of a drug.

Figure 1 highlights that unintended drug effects can occur at three different points during the drug and MR pathways. This means the effects can be considered as three different types: drug specific effects; mechanism effects; and biomarker effects. These effects are best presented in terms of the statin example discussed previously. Drug specific effects are effects that relate only to the particular drug received - this means different statins produce different effects. For example, it has been suggested that there is an increased risk of fatal rhabdomyolysis associated with cerivastatin compared with other statins and this has led to them being withdrawn from the market. (14–18) Mechanism effects are effects produced by enzyme manipulation but not biomarker changes i.e. manipulation of HMGCR inhibitor but not changes in LDL cholesterol. (19–21) For example, statins may have lipid independent effects, such as improvement of endothelial function, though there is limited direct evidence for this in humans at present. (21–23) Finally biomarker effects are the effects that result from changes in the biomarker, i.e. changes in LDL cholesterol level, which occur regardless of the mechanism used to induce that change. For example reduced LDL cholesterol appears to increase the risk of type 2 diabetes independent of the mechanism of LDL reduction. (11,24–27) Understanding the differences between these effects is important as it is key to understanding what is possible with MR in this context.

The potential of MR for predicting unintended drug effects in the future is highlighted by studies that have assessed recognized unintended drug effects. Consider once more the example of statins prescribed for the prevention of CHD. Statins increase the risk of new-onset type 2 diabetes – a risk that is recognized by both the Medicines and Healthcare products Regulatory Agency (MHRA) in the UK and the Food and Drug Administration (FDA) in the US. (28,29) This risk was originally assessed using evidence from a meta-analysis of randomized statin trials. (25) Since the recognition of this adverse drug event, Swerdlow et al conducted a MR study to assess whether the increase in new-onset type 2 diabetes risk is a result of the inhibition of HMGCR i.e. the enzyme targeted by statins. To do this, they used the SNP rs17238484 as a proxy because it is located on the HMGCR gene and has been associated with lower LDL cholesterol in a large genome-wide study of lipids. (24,30) Swerdlow et al found “each additional rs17238484-G allele was associated with a mean 0·06 mmol/L (95% CI 0·05–0·07) lower LDL cholesterol and higher body weight (0·30 kg, 0·18–0·43), waist circumference (0·32 cm, 0·16–0·47), plasma insulin concentration (1·62%, 0·53–2·72), and plasma glucose concentration (0·23%, 0·02–0·44).” (11) This led them to conclude that inhibition of HMGCR “at least partially” explains the increased risk of type 2 diabetes. In principle MR could potentially have provided evidence of this effect prior to licensing and before the exposure of large numbers of patients. Further opportunities to predict unintended drug effects are detailed in Table 1.

**Table 1:** Opportunities to predict unintended drug effects using MR with potential genetic variants identified from the GTEx eQTL catalog (69) using MR-Base (40).

| Drug | Potential proxy genetic variant(s) | Mechanism | Biomarker | Target disease |
| --- | --- | --- | --- | --- |
| Aldehyde dehydrogenase inhibitors | rs201649047<br>rs11066055<br>rs592967<br>rs11608345<br>rs111900779<br>rs7963329<br>rs57186456<br>rs201574057<br>rs847892<br>rs200037659<br>rs11066018 | Acetaldehyde dehydrogenase | Acetaldehyde | Alcohol dependence |
| Angiotensin-converting enzyme inhibitors | rs4311<br>rs6504163<br>rs4277405<br>rs4330 | Angiotensin-converting enzyme | Blood pressure | Hypertension |
| Carbonic anhydrase II inhibitors | rs11329721<br>rs10090196<br>rs3839863<br>rs13282987<br>rs62512073<br>rs79597773 | Carbonic anhydrase II | Intraocular pressure | Open-angle glaucoma |
| Cholesterylester transfer protein inhibitors | rs821840<br>rs11508026<br>rs201940645 | Cholesteryl ester transfer protein | Low density lipoprotein cholesterol | Coronary heart disease |
| Ezetimibe | rs411279633<br>rs199683176<br>rs217402<br>rs11972520<br>rs745833 | Niemann-pick C1-like 1 | Low density lipoprotein cholesterol | Coronary heart disease |
| Fatty acid amide hydrolase inhibitors | rs7520850<br>rs6429600<br>rs2145409<br>rs7555240<br>rs2145409<br>rs2145409<br>rs6429600<br>rs56083025<br>rs35361357<br>rs56083025<br>rs6429600<br>rs7555240<br>rs4660346<br>rs2145409<br>rs56083025<br>rs11804189<br>rs12217016<br>rs201127808 | Fatty acid amide hydrolase | Anandamide | Inflammatory chronic pain |
| Gonadotrophin-releasing hormone antagonists | rs28526365<br>rs12651577<br>rs145250522<br>rs17634475<br>rs12651577<br>rs141552662<br>rs147425774<br>rs398107462<br>rs145250522<br>rs11283415<br>rs199604647<br>rs147425774<br>rs1484186<br>rs71219068<br>rs13124793<br>rs11282189 | Gonadotropin-releasing hormone receptors | Luteinising hormone | Prostate cancer |
| Proprotein convertase subtilisin/ kexin type 9 inhibitors | rs2495503<br>rs34232196<br>rs479910 | Proprotein convertase subtilisin/kexin type 9 | Low density lipoprotein cholesterol | Coronary heart disease |
| Selenium | rs673752<br>rs28326<br>rs7714738<br>rs7356546<br>rs146701923<br>rs72764983<br>rs248381<br>rs485851<br>rs6453427<br>rs684277<br>rs1717567<br>rs1274984<br>rs7719892 | Dimethylglycine<br>dehydrogenase | Plasma<br>selenium | Prostate cancer |
| Statins | rs17244897 | 3-hydroxy-3-methylglutaryl-<br>coenzyme A<br>reductase | Low density<br>lipoprotein<br>cholesterol | Coronary heart<br>disease |

*MR: Mendelian Randomization. Note: Several of these drugs have already been the subject of MR studies, including ezetimibe and statins. However, these drugs could still benefit from further research, particularly combining MR with a ‘phenotype screen’ (MR-PheWAS) in order to generate hypotheses. (11,26,37,70)*

The use of genetics for pharmacovigilance and drug repurposing has previously been discussed; however, the potential of MR for this purpose is yet to be fully realized. (31,32) Until now the discussion has focused on the use of genome- and phenome-wide association studies (GWAS and PheWAS respectively). (33–36) GWAS search for the genetic variants associated with a given phenotype while PheWAS search for phenotypes associated with a given genetic variant. In these studies, the genetic variant will be a proxy for the exposure and the phenotype will be an unintended drug effect. MR extends the use of genetics for pharmacovigilance and drug repurposing as it can either be used on a single outcome, or combined with a ‘phenotype screen’ for the prediction of effects of drugs on a wide range of outcomes. The concept of MR with phenotype screening was first introduced by Millard et al, who proposed MR-PheWAS. MR-PheWAS uses “automated screening with genotypic instruments to screen for causal associations amongst any number of phenotypic outcomes”. (37) This approach is hypothesis-free and could therefore be of great use for generating hypotheses concerning potential unintended drug effects, particularly pre-approval of a drug. Limited phenotypic screening with MR has previously been demonstrated in the literature by the Interleukin 1 Genetics Consortium in their investigation of the long-term effects of interleukin 1 (IL-1) inhibition. (38) This study used a GWAS in order to inform the construction of a genetic score. The score combined the information for two SNPs, rs6743376 and rs11687782, which were upstream of the 1L1RN gene and had been shown in the GWAS to be independently associated with circulating IL-1 receptor antagonist concentration. (39) Since this study, the development of databases of harmonized summary GWAS results, such as MR-Base (http://www.mrbase.org/), has made the implementation of MR in this way much simpler. (40) The use of MR therefore has immense potential for the prediction of adverse drug events and drug repurposing opportunities, with or without a priori hypotheses. We will now discuss several advantages of using this method.

One of the key benefits of MR is that it can be used to predict drug effects that are unlikely to be affected by confounding by indication. Confounding by indication occurs in observational studies when the factors predisposing a patient to receive treatment are also the factors related to an increased risk of experiencing an outcome. (41) This can induce an artificial association between the drug exposure and an observed outcome. MR minimizes confounding by indication because the genetic variant used to proxy drug exposure is unlikely to be affected by the indications for such drug exposure. Let us continue with the example of statins prescribed for the prevention of CHD. Existing cardiovascular disease is a major indication for taking statins. At the same time, patients with cardiovascular disease are also at increased risk of death. This can induce an observational association between statin use and increased risk of cardiovascular death. But this association is not caused by statins, it is due to the indication: risk of cardiovascular disease. (42) MR reduces this confounding as the SNP located on the HMGCR gene, used to proxy exposure to statins, is a germline variant so will not change as a result of the indication i.e. cardiovascular disease.

In addition to addressing confounding by indication, MR is potentially more robust to confounding by environmental and lifestyle factors and reverse causation. As highlighted previously, this is because MR uses germline genetic variants that generally cannot be affected by environmental and lifestyle factors later in life. (43,44) Consequently, if a genetic variant is associated with an outcome through its association with a drug effect (and no other pathway), it is likely to be because the genetic variant causes the outcome. (45) Thus, MR can provide robust evidence about the causal effects of intervening on specific biological pathways. This is particularly important when considering physiological factors that change over the life course, such as fibrinogen concentration and LDL cholesterol, because the association of such factors with the outcome is likely to be heavily confounded by environment and lifestyle factors, as well as potentially being subject to reverse causation. For example, studies have previously found that people with higher fibrinogen concentrations have a higher risk of CHD, yet people with genetically higher fibrinogen concentrations do not appear to have this higher risk. This has led to questions about the causality of the relationship. (46–52) MR studies have since found that lowering fibrinogen concentration is unlikely to reduce CHD. (53,54) These results should not be unaffected by reverse causation, which suggests that interventions that lower fibrinogen concentration are not suitable drug candidates.

MR can also be used for the prediction of unintended drug effects either pre- or post-approval of a drug, in a non-experimental setting. For example, consider the potential use of selenium dietary supplements for the prevention of prostate cancer. The Selenium and Vitamin E Cancer Prevention Trial (SELECT) found that selenium did not lower prostate cancer risk but did increase the risk of type 2 diabetes. A MR study, conducted after the trial, found genetically elevated selenium was not associated with prostate cancer risk and was positively associated with type 2 diabetes risk (Martin RM, personal communication). Implementation of MR prior to the trial could therefore have been an informative step in the assessment of selenium as a possible chemoprevention target. Investigation of drug effects pre-approval is particularly appealing in relation to adverse drug events. Even without a hypothesis for a potential unintended drug effect, the MR-PheWAS approach discussed earlier can be used in order to generate hypotheses. The appeal of using MR prior to a trial is that it can be used to pre-specify likely adverse outcomes to be measured during the trial and determine major unintended drug effects that will prevent market authorization. Conducting MR prior to clinical trials could therefore help reduce the possibility of exposing trial participants to unnecessary risks and harm and, ultimately, the general population at the point of approval if the drug obtains market authorization. It also has the potential to increase the efficiency of drug development, saving both time and money in this process.

A further advantage of MR is that it can be used to predict the combined effects of drugs. This is important as many medicines are only licensed for use when other treatments are either being used concurrently or have been previously used and failed. For example, trials are ongoing for proprotein convertase subtilisin–kexin type 9 (PCSK9) inhibitors to reduce LDL cholesterol. As previously stated, statins are currently used for this purpose and have been associated with an increased risk of type 2 diabetes. Consequently, there have been concerns that PCSK9 inhibitors may have similar effects. This has led to several MR studies being conducted in order to anticipate the effect of PCSK9 inhibitors on type 2 diabetes risk. (26,27,55) These include a study by Ference et al, which compared the risk of cardiovascular events and type 2 diabetes due to variation in PCSK9, HMGCR or both. Hence allowing “inferences about the potential clinical benefit and safety of treatment with a PCSK9 inhibitor as compared with treatment with a statin” to be made. (27) The study found PCSK9 variants to have a similar effect as HMGCR variants on both the risk of cardiovascular events (OR 0.81, 0.74-0.89 vs OR 0.81, 0.72-0.90) and the risk of type 2 diabetes (OR 1.11, 1.04-1.19 vs 1.13, 1.06-1.20) for each 10 mg per deciliter decrease in LDL cholesterol level. The similarity of the variant effects and additional analyses, which assessed potential shared pathways, led Ference et al to suggest that the cause of increased type 2 diabetes risk may be related to a LDL receptor-mediated pathway, i.e. may be a biomarker effect. This indicates that switching patients from statins to PCSK9 inhibitors because of type 2 diabetes risk may not be beneficial. Ference et al also found the combination of PCSK9 and HMGCR variants to be “independent and additive”. This suggests prescription of these drugs simultaneously will both amplify the benefits of these drugs and increase their associated type 2 diabetes risk. The notion of ‘additive’ effect is particularly important here as trials of PCSK9 inhibitors are currently being conducted against a background of taking statins, making predicting trial results using other methods more difficult. Overall, the results of the study suggest that PCSK9 inhibitors are likely to offer similar clinical benefits and risks as statins and will have an additive effect if prescribed together. The use of multiple lipid-lowering therapies is also considered by Robinson et al, who use pooled trial data to assess the safety of very low LDL cholesterol levels when using the PCSK9 inhibitor alirocumab. Consideration of very low LDL cholesterol levels is important as the use of concurrent treatments grows. This study concluded that “LDLC levels < 25 or < 15 mg/dl on alirocumab were not associated with an increase in overall treatment emergent adverse event rates or neurocognitive events, although cataract incidence appeared to be increased in the group achieving LDL-C levels < 25 mg/dl”. (56) MR could be used in this situation to further investigate cataracts as a potential unintended drug effect. Conclusions obtained from MR studies, such as those presented by Ference et al, are valuable as they provide information about concurrent use of drugs, which is common in clinical practice, and can help inform whether a drug is suitable for a patient for whom other drugs have failed. The example from Robinson et al highlights the opportunity to triangulate evidence from trials with an MR study.

Additionally, the use of genetic variants to proxy an exposure in MR can address missing or incomplete exposure, outcome or confounder data. (57) GWAS are increasingly publishing the associations between all genetic variants and their outcome. This means the associations between a genetic variant and an outcome can be looked up in databases of GWAS results (such as MR Base, http://www.mrbase.org/) and the need for new analysis of individual level data is removed. (40) Provided there are robust genetic variants for the drug exposure of interest (see Table 1 for examples) and there are large genetic association studies of the outcome, MR will not be limited by missing or incomplete data. Furthermore, MR and the related genetic methods can be used with non-genetic approaches in order to better explore the relationship between the genome and phenome. (58) Bush et al discuss how genetic data can be linked with data from electronic health records and epidemiological studies in order to better characterize “the impact of one or more genetic variants on the phenome” in the PheWAS setting. (59) An MR-PheWAS that implemented such an approach could therefore be an even more powerful tool for the prediction of unintended drug effects. Alternatively, germline genetic variants can be used to proxy drug activity in studies that relate to potential unintended drug effects in patient groups, without having to implement a full-scale PheWAS. Walley et al demonstrate this by using healthy patients genotyped for PCSK9 loss of function SNPs to test their hypothesis that PCSK9 inhibition is involved in pathogen lipid clearance and could improve septic shock outcomes. The use of healthy patients for this study allows investigators to avoid “the heterogeneity of septic shock” that may bias their results. The combination of MR with this type of study could be used to enhance evidence for unintended drug effects in specified patient groups in the future. (60)

Finally, MR can help to differentiate between unintended drug effects in a way that is difficult with other approaches. Earlier we highlighted that unintended drug effects can be considered as three different types: drug specific effects; mechanism effects; and biomarker effects. MR can be used to distinguish mechanism and biomarker effects. Unfortunately, you are unlikely to find genetic variants that proxy specific drugs and so it is difficult to distinguish drug specific effects from other effect types. Mechanism effects can be distinguished using MR if there is evidence of displacement of genetic variants that proxy one mechanism, which effects a downstream phenotype, from the variants that proxy an alternative mechanism, which effects the same downstream phenotype. This is illustrated in Figure 2. For example, a difference between PCSK9 and HMGCR variants, given their effect on LDL cholesterol, would suggest a mechanism effect. This approach could be implemented by assessing the displacement from MR-Egger of particular variants using a measure such as Cook’s distance. This has previously been demonstrated in the literature by Corbin et al, who conducted a study to resolve the relationship between body mass index (BMI) and type 2 diabetes. The study found that pleiotropy (when a genetic variant influences multiple phenotypes that are thought to be distinct) may be present with respect to the outcome, type 2 diabetes, for the genetic variant rs7903146, indicating that it “influences type 2 diabetes through an alternative pathway (other than BMI)”. This method could easily be applied to the assessment of a pharmacological intervention in a similar way.

**Figure 2:**
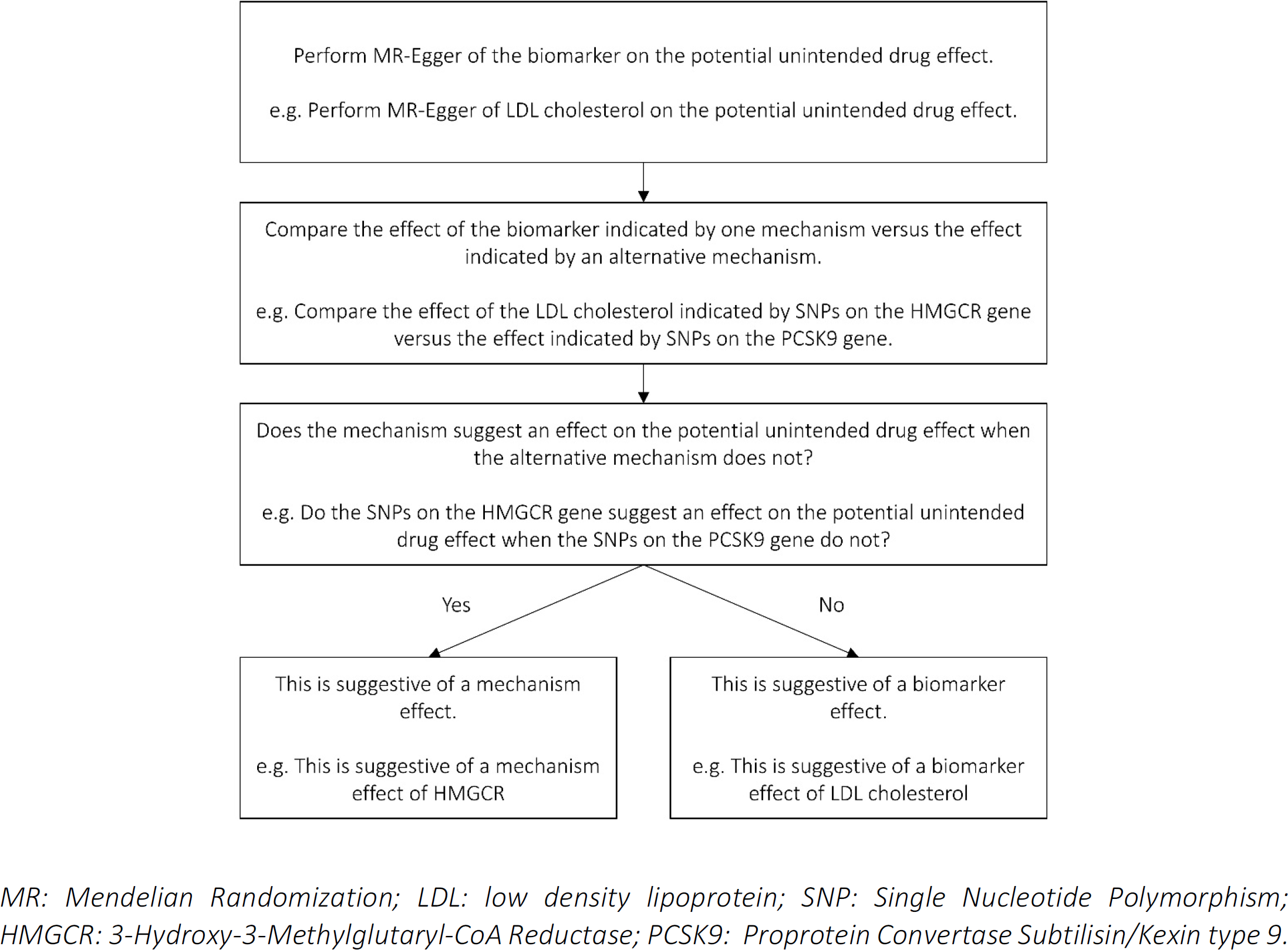
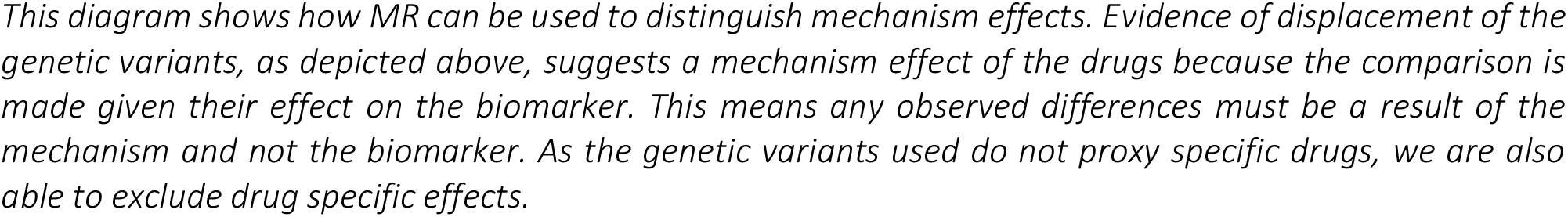
The process by which MR can be used to distinguish mechanism effects of drugs.

Biomarker effects may also be distinguished by MR as it enables a formal statistical comparison of the effect of a biomarker influenced by different drug-related mechanisms to be made. (26,61,62) For example, as described earlier, Ference et al suggested that the cause of increased diabetes risk may be related to a LDL receptor-mediated pathway as this risk is associated with both the variants for HMGCR, the mechanism of statins, and the variants for PCSK9, the mechanism of PCSK9 inhibitors. (27) This indicates that increased risk of type 2 diabetes is likely to be a biomarker effect, i.e. a result of changes in LDL cholesterol level, rather than a drug specific or mechanism effect of statins. Other genetic studies have found similar results. (26,27,55) MR can therefore distinguish mechanism and biomarker effects by using different proxies for exposure that occur at different points on the common part of the drug and MR pathways. This distinction can be difficult to achieve using the existing observational methods but is important as the effects may have different implications and therefore require different action in order to be resolved.

As with all methods, MR has limitations. In particular, the method is not suited to detecting rare unintended drug effects due to the power and data availability issues associated with such outcomes. For example, it has been suggested that rhabdomyolysis may be a mechanism effect of statins that is more pronounced for cerivastatin, rather than a drug-specific effect of cerivastatin. The global incidence of rhabdomyolysis is unknown but it is thought to be rare with an estimated 26,000 cases per year occurring in the US according to the 1995 National Hospital Discharge Survey. (63,64) This means MR studies are likely to be underpowered to detect rhabdomyolysis as a mechanism effect of statins. (65–68) In addition to this, unintended drug effects may be missed due to the choice of genetic variant used in the MR study. This can happen if you chose a genetic variant to proxy exposure downstream of the effect you are interested in. For example, if you chose a genetic variant related to LDL cholesterol level, i.e. at the biomarker level, to investigate statins then the mechanism effects, i.e. the lipid-independent effects such as improved endothelial function, will be missed. In addition to this, genetic association studies often investigate only common genetic variants or combine the effect of rare genetic variants. This results in a situation where individual genetic variants may explain very little of the observed variation. Careful consideration must therefore be given to the choice of genetic variant when conducting an MR study.

MR offers a novel and appealing approach for the prediction of unintended drug effects. The method can overcome some of the limitations associated with existing methods in this field, as well as provide additional benefits, such as use pre-approval of a drug. We encourage the use of MR to triangulate evidence of unintended drug effects and enhance existing methods, such as in silico approaches. The potential of MR as a pharmacovigilance and drug repurposing tool is yet to be realized. Future use of this method in the development of drug profiles could both help prevent adverse drug events and identify novel indications for existing drugs.

